# T-lex2: genotyping, frequency estimation and re-annotation of transposable elements using single or pooled next-generation sequencing data

**DOI:** 10.1101/002964

**Authors:** Anna-Sophie Fiston-Lavier, Maite G. Barrón, Dmitri A. Petrov, Josefa González

## Abstract

Transposable elements (TEs) constitute the most active, diverse and ancient component in a broad range of genomes. Complete understanding of genome function and evolution cannot be achieved without a thorough understanding of TE impact and biology. However, in-depth analysis of TEs still represents a challenge due to the repetitive nature of these genomic entities. In this work, we present a broadly applicable and flexible tool: T-lex2. T-lex2 is the only available software that allows routine, automatic, and accurate genotyping of individual TE insertions and estimation of their population frequencies both using individual strain and pooled next-generation sequencing (NGS) data. Furthermore, T-lex2 also assesses the quality of the calls allowing the identification of miss-annotated TEs and providing the necessary information to re-annotate them. The flexible and customizable design of T-lex2 allows running it in any genome and for any type of TE insertion. Here, we tested the fidelity of T-lex2 using the fly and human genomes. Overall, T-lex2 represents a significant improvement in our ability to analyze the contribution of TEs to genome function and evolution as well as learning about the biology of TEs. T-lex2 is freely available online at http://sourceforge.net/projects/tlex/.

**Abbreviations:**

TE: transposable element
NGS: next-generation sequencing
LTR: long-terminal repeat
TSD: target site duplication
PTS: putative target site
PE: paired-end

## INTRODUCTION

The key question in genomics is how genomes vary and evolve at both large and fine scales. In order to answer such question, we need to be able to study genome variation, *i.e.* identifying and analyzing functionally relevant SNPs (Single Nucleotide Polymorphisms) and structural variants, both within and between populations. Next-generation sequencing (NGS) technology has revolutionized this field by allowing one to study variation of a large number of individuals and even cells within individuals (1–3). Unfortunately, NGS technology generally provides fairly short, even if extremely abundant, sequencing reads and thus is not perfectly designed for the study of structural variants (4).

The study of one class of structural variants, transposable elements (TEs), is particularly difficult to carry out using NGS data. TEs are repetitive, ubiquitous and dynamic components of genomes that often vary in genomic location among members of the same species. TEs are classified in three orders: DNA transposons, long-terminal repeat (LTR) elements, and non-LTR elements, and each TE order is represented by several TE families (5,6). This classification highlights the biological diversity in terms of sequence, dynamics and evolution of these repetitive elements that makes their analysis challenging.

TEs represent a large part of most of the eukaryotes genomes (7). A recent study suggests that more than two-thirds of the human genome is composed of TEs (8), and in plants, TEs may represent up to 90% (9). TEs are responsible for a large number of mutations both in populations and somatically within individuals (10,11). Although most of the TE-induced mutations are deleterious, evidence for an adaptive role of TE-induced mutations is starting to accumulate (12–16). TEs can provide active promoter, splice site, or terminator features that can affect the expression, structure and function of nearby genes (17). TEs are also involved in the creation of transcriptional regulatory networks, and in the generation of chromosomal rearrangements (12,18). Thus, knowing the role of TEs on genome dynamics and evolution, it is crucial to identify and quantify their impact (19,20). Given the increased number of sequenced genomes, a tool that allows routine and automatic genotyping of individual TE insertions using a large number of NGS samples is needed.

Recently several accurate computational approaches have been developed to analyze TE insertions using NGS data (10,21-25). Designed for specific projects, all these tools were built with a limited number of features, *e.g.* some tools were designed to specifically call particular families of TEs (10), while others were designed exclusively for pooled NGS data (25). Unfortunately, none of the currently available tools allows a complete and in-depth analysis of individual annotated TE insertions.

With the idea of providing a tool to automatically call TE insertions in multiple genomes and to estimate their population frequency, we recently designed an integrated, flexible pipeline called T-lex (26). While most of the available approaches only detect the presence of TE insertions, T-lex launches two complementary and distinct TE detection modules: one to detect the “presence” and another to detect the “absence” of a TE insertion. Both detection approaches are based on the analysis of the junction sequences of a TE insertion and its flanking sequences as defined by genome annotation. Thus, the accuracy of the TE calls mainly depends on the quality of the annotation and more specifically on the delimitation of the individual TE insertions. Unfortunately, the detection of the precise TE insertion sites is still one of the main challenges of the TE annotation process (27). The veracity of the TE calls also depends on the genomic environment of the TE insertion. For instance, a TE insertion located inside a duplication with one of the copies lacking the TE insertion can be miscalled as heterozygote (*i.e.,* both present and absent). In addition, a TE insertion flanked by other repeats such as low-complexity regions or other TE sequences can also be miscalled.

Here, we present T-lex2, the version 2 of the original T-lex pipeline (26). Besides improving the accuracy of the TE frequency estimate, T-lex2 now assesses the quality of the TE calls. To achieve this, T-lex2 uses information from (i) a new module specifically designed to identify target site duplications (TSDs), and (ii) the genomic context of each insertion, to identify putatively miss-annotated TE insertions highly likely to produce wrong calls. TSDs are short duplications, from two base pairs to 20 bp, flanking the TE sequences as a result of the transposition mechanism of most TE families. While current TSD detection approaches require knowing the biology of the TEs and are limited to LTR or non-LTR elements, T-lex2 TSD detection module works without *a priori* knowledge, and for all type of TEs. This new module allows precisely delimiting the location of each TE insertion. In this new version, frequency of TE insertions can be estimated using individual and pooled NGS data.

We provide evidence that the new features of T-lex2 help improving the genotyping and frequency estimate of TE insertions, and allow re-annotating TE locations using both individual and pooled *Drosophila melanogaster* genomic data. We also demonstrate that T-lex2 provides accurate TE calls in genomes other than that of *D. melanogaster* by analyzing TE insertions.

## MATERIAL AND METHODS

### T-lex2 pipeline overview

T-lex2 is composed of five modules that can be run with individual strain or pooled NGS data (Figure 1). (i) The TE-analysis module investigates the flanking regions of the known TE insertions (*i.e.* annotated in a reference genome) to identify those likely to return wrong calls. (ii) The TE-presence detection and (iii) the TE-absence detection modules combine mapping and read depth coverage information to identify reads providing evidence for the presence and for the absence, respectively, of the known TE insertions. (iv) Then the TE-combine module combines the results of the presence and absence detection modules to genotype TEs and/or to estimate their frequencies in populations. Finally, (v) the TE-TSD detection module annotates TSDs in an unbiased and accurate way. Each of these modules can be launched independently and provides a large number of options allowing the user to carry out flexible and customizable analyses. A detailed manual describing step-by-step how to run T-lex2 and listing all T-lex2 options is available on line at http://petrov.stanford.edu/cgi-bin/Tlex2_manual.html.

**Figure 1.**
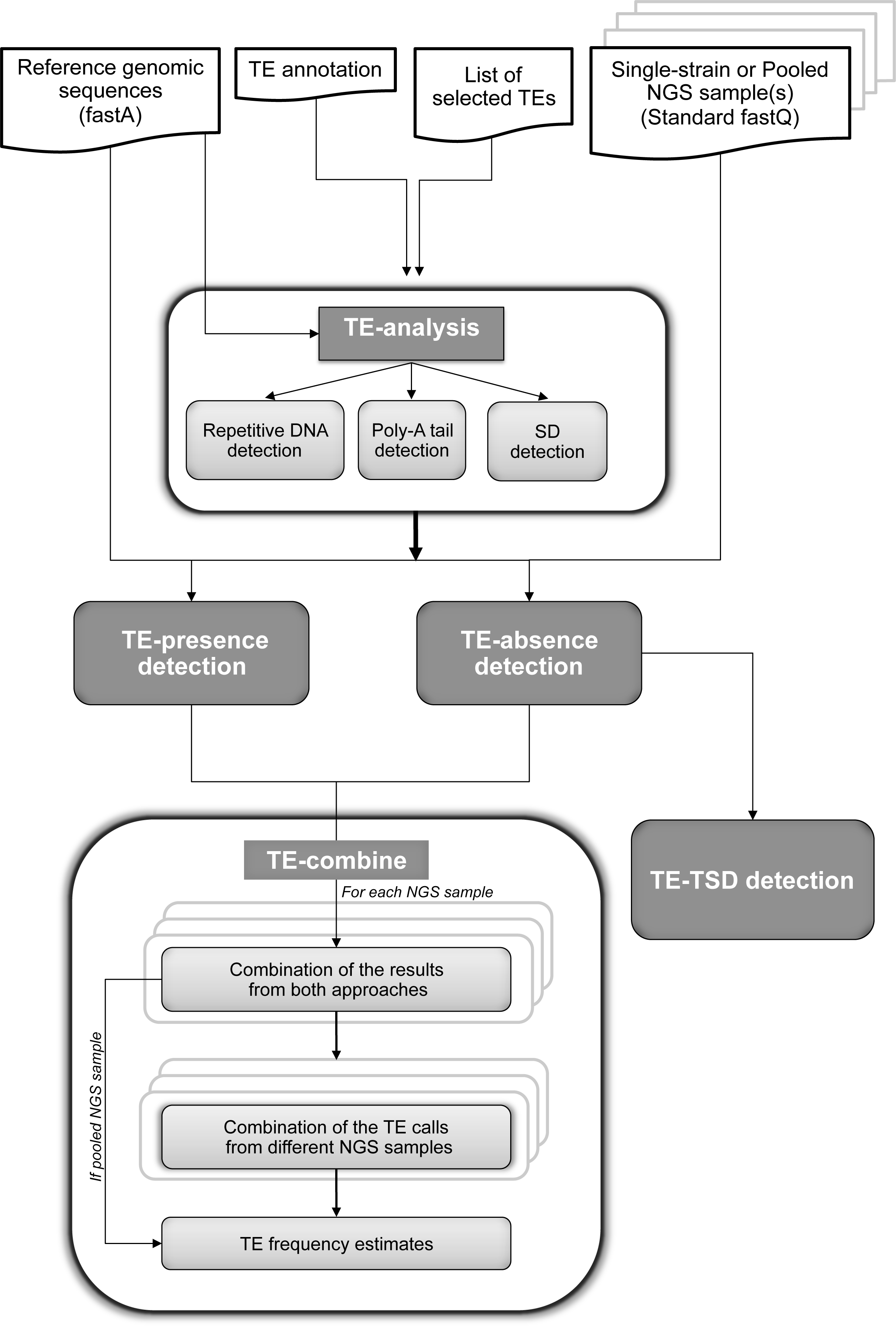
T-lex2 pipeline. Schematic representation of the five different T-lex modules (dark gray boxes) and of the input files (white boxes) required to run T-lex2. SD: Segmental Duplication.

**TE-analysis module**. This module analyses the flanking sequences of each known TE insertion to identify features that might interfere with the presence and absence detection modules. The TE-analysis module (i) provides information about the presence of repetitive elements using RepeatMasker (28), (ii) identifies miss-annotated poly(A) tails, and (iii) flags the TE insertions that are part of segmental duplications.

i. To identify the presence of repetitive elements, the TE-analysis module extracts the flanking sequences of each TE insertion (125 bp by default; see option “-f”) and launches RepeatMasker to annotate the TEs, simple repeats and low-complexity regions (28). The option “-s” allows specification of the name of the model organism and thus specifies the repeat library that needs to be used by RepeatMasker (28). If at least one flanking region shows a repeat density greater than the pre-specified value (50% by default; option “-d”) the TE insertion is flagged as flanked by repetitive elements. By default, these TE insertions are filtered out. This filtering step can be bypassed using the option “-noFilterTE”.
ii. A new feature of the TE-analysis module identifies putatively miss-annotated poly(A) tails by searching for stretches of A’s or T's (by default more than five base pairs) at the TE junctions, without *a priori* knowledge of the TE type. Non-LTR elements harbor poly(A) tails at their 3'end that are known to be highly variable in length. Such sequences are therefore very difficult to annotate automatically. As RepeatMasker only annotates repeats longer than 20 bp, it cannot be used to identify short stretches of A's or T's that may correspond to miss-annotated poly(A) tail sequences (28).
iii. When a TE insertion is part of duplication, other copies of the same duplication may not contain the insertion. The analysis of the flanking sequences of such TE insertions may lead to call this TE insertion as present and absent while it should only be called as present. We added one new feature in the analysis of the TE flanking regions to identify the TE insertions fully or partially part of segmental duplications. For each TE insertion, the two extracted flanking sequences are blated against a reference genome (29) (BLAT v. 34 default parameters; http://www.soe.ucsc.edu/∼kent). When more than 50% of a flanking sequence matches at more than one location on the reference genome with more than 90% of sequence identity, the TE insertion is flagged as part of a duplication. TE insertions partially part of duplication (notified as “sd_left” or “sd_right”) are distinguished from the TE fully part of a duplication (notified as “sd”). When a TE is fully part of a duplication while another copies do not contain the TE insertion, this TE insertion is notified as “sd_noTE”.

The analysis of the TE flanking sequences including the RepeatMasker output, whether the TE is flanked by a longer than annotated poly(A) tail, and the segmental duplication analysis results, are stored in a sub-directory called “Tanalyses”.

**TE-presence detection module**. The presence detection module detects sequencing reads overlapping the flanking regions of the TE insertion (Figure 2A). The junction sequences of each TE insertion are extracted and used as reference sequences. By default, each TE junction sequence encompasses the terminal TE sequence (60 bp from each end) and the flanking sequence (1 kb on each side). The lengths of the TE sequence and the flanking sequence can be changed using the options “-b” and “-j” respectively. The reads are mapped on these new extracted reference sequences (Figure 2A) and are used to build a contig using Phrap (30) (version 1.090518; http://phrap.org). T-lex2 uses only the reads mapping with a minimum Phrep quality score (30 by default; this can be changed using the option “-minQ”). The pipeline also reports the number of reads mapped on the TE junction at the minimum Phrep quality score (Figure 2A). Additionally, T-lex2 also requires a minimum length of the match inside and outside the TE sequence, and a minimum read sequence identity (Figure 2A). These two parameters are set at 15 bp and 95% by default and can be modified using the options “-limp” and “-id”, respectively. By default, multiple alignments of the contigs for each TE side are also returned. The analysis and TE calls from this module are stored in a sub-directory called “Tpresence”.

**Figure 2.**
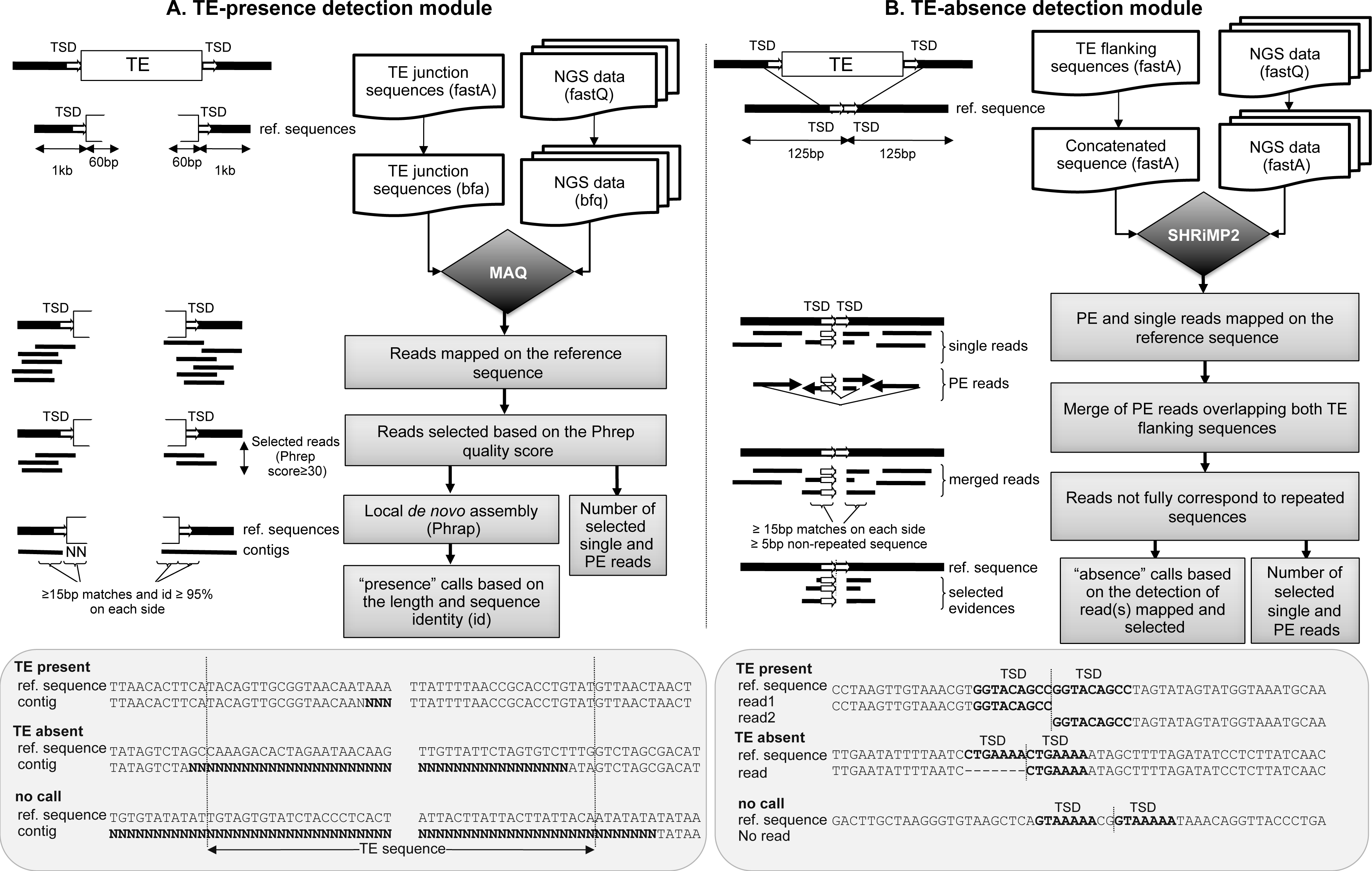
T-lex2 TE-presence detection and TE-absence detection modules. (A) TE-presence detection module is based on the mapping of the NGS reads on the TE insertion junctions. (B) TE-absence detection module is based on the mapping of NGS reads on the putative ancestral genomic sequence prior the TE insertion. Input files required to run each module (white boxes), the different steps of the pipeline (light grey boxes), and examples of reads providing evidence for the presence, absence, and no call reads are depicted.

**TE-absence detection module.** The absence module detects reads overlapping both sides of the TE insertion, *i.e.* reads spanning across the TE insertion site (Figure 2B). It starts by extracting the two flanking sequences for each TE insertion (125 bp each by default; see option “–f”). The two sequences are then concatenated and the new constructed sequence is used as the reference sequence for the absence detection (Figure 2B). This new constructed sequence approximates the ancestral sequence prior to the TE insertion. However, note that traces of the TE insertion mechanism, called TSDs, may also be encompassed in the new constructed sequence and located at the TE insertion breakpoint. The length of the extracted sequences should be longer than the reads themselves and total length of the new constructed sequence should be similar to the insert size of the paired-end (PE) data in order to get reads and/or pairs spanning the TE insertion breakpoints. T-lex2 uses now by default a length of 125 bp matches for libraries of 100 bp read length and an insert size of 250 bp. The reads are then mapped on the new constructed sequence using SHRiMP2 (31) (version 2.2.1, October 2011), the new version of SHRiMP that can now handle PE read sequencing data. SHRiMP2, as SHRiMP, was specifically designed to handle long gaps and polymorphisms (31,32). Such features allow the mapping of reads despite the presence of the TSDs or despite miss-annotation. To handle long gaps, SHRiMP2 is launched with a Smith-Waterman gap open score of −40 and a gap extension score of −1 for both query and reference sequences. We also allow higher divergence with a Smith-Waterman mismatch score of −20. SHRiMP2 maps the reads in every pair together (31) (see option “-pairends”). If a pair does not map, SHRiMP2 attempts to map each read individually (31). If the two reads of a pair map to the new constructed reference sequence, only the pairs with reads mapping on opposite strands are selected (*i.e.,* concordant pairs). When the two reads from the pair overlap in the mapping, the absence module merges them and considers the pair as a single read. The absence module specifically looks for “reads” spanning the TE junction with at least 15 bp (parameters by default; see option “–v”) of overlap on each side. T-lex2 then runs RepeatMasker on the selected reads (see option “-noint”) in order to test whether the mapping is due primarily to simple repeats or low-complexity regions (28). After this step, T-lex2 selects the reads that have at least five non-repetitive (low-complexity or satellites) base pairs at each end that both map to the flanking regions. Because of the stringency of our approach, even a single read mapping to both flanking sides and thereby spanning the insertion site of the TE is sufficient for the absence detection module to classify the TE insertion as “absent” (Figure 2B). If no reads overlap the TE junction, two possibilities are considered: (i) the TE is present in the strain (or fixed in the pool sample) or (ii) the coverage of the data used is insufficient to detect reads providing evidence for the absence. In order to distinguish between these two possibilities, the absence detection module checks whether reads map on both flanking sides but fail to map over the junction. If this is the case, given that the flanking regions are similar in length to the reads themselves, the module concludes that the TE is present.

If reads do not map to both flanking sides and do not map over the junction site, the module concludes that the coverage is insufficient and returns a “no call” as a result. By default, the multiple alignments of the contigs for each TE breakpoint are also returned. The analysis and TE calls from this module are stored in a sub-directory called “Tabsence”.

**TE-combine module**. The results from the presence and absence detection approaches can then be combined to return definitive TE detection calls. We defined five TE call categories based on the evidence for presence and absence. Briefly, a TE is called “present” or “absent” when the calls from both detection modules are congruent. A TE is called as “polymorphic” when this TE is clearly detected as present and absent (*i.e.,* the presence detection module detects it as present and the absence detection module detects it as absent). A TE is called as “present/polymorphic” when the presence detection module detects it as present while the absence detection module fails to return a call. A TE is called as “absent/polymorphic” if the TE is detected as absent by the absence detection approach, while the presence detection fails (see Table 1 in (26)).

**Table 1.** T-lex2 is an improved and expanded version of T-lex. Comparison of the original T-lex version and T-lex2 modules and features

| **T-lex2 Modules** | **Features** | **T-lex (26)** | **T-lex2** |
| --- | --- | --- | --- |
| **TE-analysis** | Detection of low-complexity regions | ✓ | ✓ |
|  | Detection of repeat-rich regions | ✓ | ✓ |
|  | Detection of terminal poly(A) stretches | ✗ | ✓ |
|  | Detection of TEs part of segmental duplications | ✗ | ✓ |
| **TE-presence detection** | Analyzes inside the TE | ✓ (20bp) | ✓ (60bp) |
| | Read quality filtered | ✗ | ✓ (phrep score $\geq 30$) |
|  | Read depth reported | ✗ | ✓ |
|  | Min. match length | ✓ (5bp) | ✓ (15bp) |
|  | Min. sequence identity | ✗ | ✓ (95%) |
| **TE-absence detection** | Analyzes on each side of the TE | ✓ (100bp) | ✓ (125bp) |
|  | PE data analysis | ✗ | ✓ |
|  | Read quality filtered | ✗ | ✓ (30) |
|  | Read depth reported | ✗ | ✓ |
| **TE-combine** | Take into account the no calls | ✗ | ✓ |
|  | Estimates using pooled sample | ✗ | ✓ |
| **TE-TSD** | TSD detection | ✗ | ✓ |
|  | Flag of putative miss-annotated TEs | ✗ | ✓ |

T-lex2 can also combine the TE calls from several NGS data from the same population and can estimate the population frequency of each annotated TE insertion (see options “–combRes” and “-combine”). To estimate the frequency, T-lex2 transforms the calls in frequencies as follows: a “present” call is associated with a 100% frequency, and absent call is associated with a zero percent frequency, a “polymorphic” call is associated with a 50% frequency (100% (present) + 0% (polymorphic)/ 2), a “present/polymorphic” call is associated with a 75% frequency (100% + 50%/ 2), and an “absent/polymorphic” call is associated with a 25% frequency (0% +50%/ 2).

Finally, T-lex2 pipeline can also estimate the TE frequency using pooled NGS data by taking into account the number of reads supporting the presence and the number of reads supporting the absence. In this case, the user just needs to add in the command line the option “-pooled”. This approach is based on two assumptions: (i) individuals in the pooled sample are equally represented, and (ii) the read-depth coverage is correlated with the occurrence of a TE sequence in a population. Based on such assumptions, we expect to observe a positive correlation between the frequency and the number of reads providing evidence for the presence. We also expect to observe a negative correlation between the frequency and the number of reads providing evidence for the absence. The number of copies of TEs from the same family may also have an impact on the number of reads mapping at the TE breakpoints suggesting that the number of reads supporting the presence can be more biased than the number of reads supporting the absence.

Taking all these expectations into account, we designed and verified a metric based on the local read depth coverage in the vicinity of the TE insertion to estimate TE frequencies from pooled data: total number of reads supporting the presence (NP) divided by the total number of reads supporting the presence (NP) and supporting the absence (NA), *i.e.* TE frequency = NP/(NP + NA).

The file combining the TE calls from both TE detection modules is called “Tresults”. The TE frequency estimates are stored in the “Tfreq” or Tfreq_pooled” if the option “-pooled” is specified.

**TE-TSD detection module**. To detect the TSD of each annotated TE insertion, this module analyzes the read alignments generated by the absence detection module (Figure 3). As all T-lex2 modules, this new module can be launched independently although it does require the read alignments from the absence detection module (see option “-tsd”). The TSD detection module looks for tandem duplications located on the absence reference sequence spanning the TE breakpoint. It starts by assembling all the selected reads supporting the absence for each TE using the Phrap program (30) (Figure 3). Because Phrap requires a minimum of three sequences to build a contig, if fewer than three reads are selected to support the absence call, the reads are considered independently (Figure 3). Each contig (or read) is then re-aligned on the absence reference sequence using BLAT v.34 program (29) (default parameters; Figure 3). Only the absence calls with a gap larger than two base pairs and located close to the TE breakpoint are selected and analyzed. If no gaps are observed, the TE-TSD detection module returns “no Gap”. If a gap is observed, the motif present in front of the gap is called the putative target site or PTS (Figure 3). Using the FastaGrep program (available from bioinfo.ut.ee/download/dl.php?file=6 and executed with the default parameters), a tool that looks for short and conserved motifs, the TSD detection module looks for the copy of the PTS on the other side of the breakpoint. Based on the sensitivity of this program and the short length of the TSDs, we only trust the matches with more than 80% sequence identity (Figure 3). T-lex2 then sorts the FastaGrep hits to identify the sequence showing the closer and highly conserved motif. If FastaGrep fails to detect a match, the TSD detection returns “noTSD” as a result, otherwise, the TSD detection module returns “detected”. In the latter case, the PTS and its closest copy sequences are also returned. Because several contigs can be generated for the same TE insertion, several TSDs can also be returned. The results of the TSD detection process are stored in a file called “TSDannot”.

**Figure 3.**
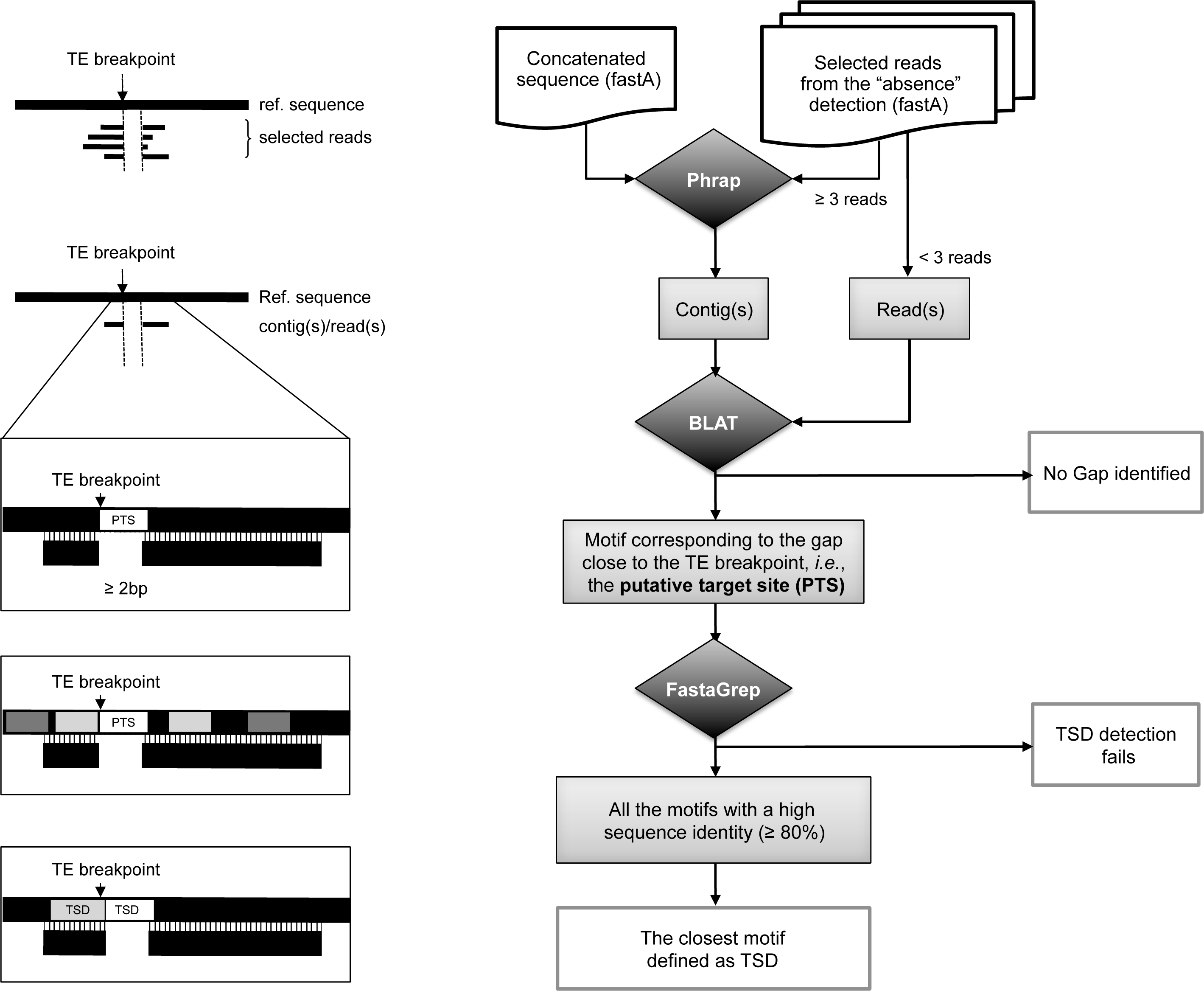
T-lex2 TE-TSD detection module. Schematic representation of the procedure to identify TSDs. Input files required to run this module and the different steps followed by the pipeline are also depicted.

TE-TSD detection module flags the TE insertions for which the TSDs cannot be clearly defined. TSD detection may fail for old TE insertions for which the TSD is too divergent, for truncated TEs, and when the boundaries of the TE insertions are not well annotated (Figure 4). The latter TEs can be re-annotated by the user after manual inspection.

**Figure 4.**
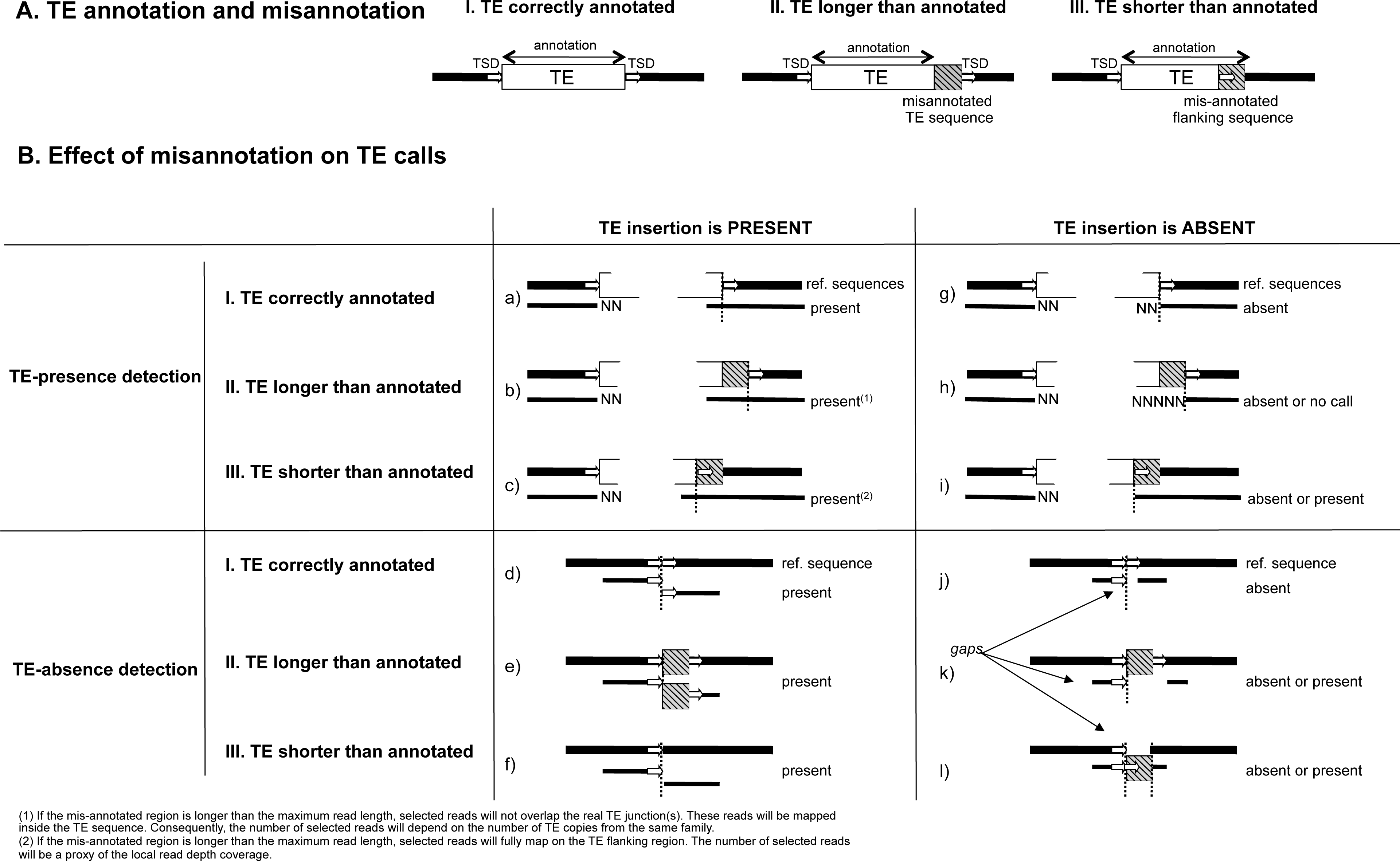
Detection of putatively mis-annotated TE insertions. (A) Schematic representation of a correctly annotated TE (I), a TE longer than annotated (II) and a TE shorter than annotated (III). (B) Effect of miss-annotation on the TE calls when the TE insertion is present and absent. When the TE insertion is absent, T-lex2 will give a correct call if the miss-annotation is short and an erroneous call in the miss-annotation is long (B.h, B.i, B.k, and B.l). (1) If the miss-annotated region is longer than the maximum read length, selected reads will not overlap the real TE junction(s). These reads will be mapped inside the TE sequence. Consequently, the number of selected reads will depend on the number of TE copies from the same family. (2) If the miss-annotated region is longer than the maximum read length, selected reads will fully map on the TE flanking region. The number of selected reads will be a proxy of the local read depth coverage.

### Re-annotation of TE insertions

Because both TE detection approaches are based on the mapping of NGS reads to the flanking TE sequences, the accuracy of the TE calls depends on having a correct TE annotation. TE-analysis and TE-TSD detection modules help identifying putatively miss-annotated TE insertions and thus putatively wrong T-lex2 calls.

A particular TE insertion can be miss-annotated because it is longer than annotated (Figure 4A.II) or shorter than annotated (Figure 4A.III) in both cases the TE-TSD detection will fail. If the TE insertion is present in a given genome, both the presence detection and the absence detection modules will provide the correct call even though the TE is longer or shorter than initially annotated (Figure 4B). However, if the TE insertion is absent, and the TE is miss-annotated the presence and the absence modules could return a wrong call (Figure 4B). For example, if the TE sequence is longer than the official annotation and the TE is absent, the gap in the reads providing the absence could be longer than the TSD size and the module could return present as a call (Figure 4B.k). On the other hand, when the TE is shorter than annotated and the TE is absent, the gap will be present in the reference sequence and not in the reads. In such situation, the module could return present as a call (Figure 4B.l).

The Tfreq_combine file of T-lex2 provides information from both the TE-analyses and TE-TSD detection modules, *i.e.* poly(A) tail longer than the official annotation and TSD detection failures, that allows classifying these TE insertions as putatively miss-annotated TEs. This information can be use to perform manual curation of these insertions and re-annotate them.

### Validation dataset

T-lex2 was validated in two datasets, one from *Drosophila melanogaster* and one from human. All the NGS data used in this work was generated using the *Illumina* technology.

Veracity of T-lex2 calls was assessed by calling a total of 755 well-studied TE insertions in *Drosophila melanogaster* strains from the Drosophila Genome Reference Panel (DGRP) project (33,34). Estimates of TE frequency based on pooled-PCR are available for these 755 TEs (34,35). TE frequency estimates based on pooled-PCR approach were categorized in four classes: “very rare” (*i.e.* TE frequency < 1.5%), “rare” (*i.e.*, TE frequency ∼2–15% but not absent), “common” (*i.e.*, TE frequency ∼10–98%) and “fixed” (*i.e.*, TE frequency > 98%) (34). The *D. melanogaster* reference genome (release 5) was downloaded from Flybase website (http://flybase.org). The annotations of the TE insertions were extracted from the release 5.43 (http://flybase.org). The resequencing data from 86 *D. melanogaster* DGRP single strains from the freeze 2 were downloaded from the DGRP website (∼20X; http://dgrp.gnets.ncsu.edu/, Supplementary Table S1). We also called the same TE insertions in the DGRP population using a high-depth sequencing data from a pool of 92 *D. melanogaster* DGRP strains (∼60X) (36). 84 strains are common in the two datasets (Supplementary Table S1). To investigate the effect of different read-depth coverage and different number of flies in pooled samples, we also used T-lex2 to analyzed TE frequencies in four other pools of lower depth coverage described in Zhu *et al.* 2012 (36).

Experimental validation of T-lex2 results was also done for 11 TE insertions known to be polymorphic (35) in 24 single *D. melanogaster* North American DGRP strains using a single PCR approach (Supplementary Table S1). The PCR conditions and primers used were as described in Gonzalez *et al.* 2008 (35).

In humans, the performance of T-lex2 was evaluated using high-depth NGS data for a CEU individual (NA12878) and the TE annotations from the Mobile Element Insertions (MEI) catalog from the 1000 Genomes Project (37). The human reference genome used was Mar 2006 NCBI36/hg18. Out of the 2,010 reference MEI reported in Steward et al (2011) (37) we analyzed 1,549 insertions that (i) have genomic coordinates with confidence intervals smaller or equal to 1 according to Steward et al (2011), (ii) could be unambiguously mapped to RepeatMasker annotations within a range of +−100 bp, and (iii) have been analyzed in the CEU trio dataset. RepeatMasker chromosome, strand and coordinates of these 1,549 TEs were used to build the TE annotation file and TE list that was used as input for T-lex2. All the TE insertions analyzed have been previously validated by PCR and/or algorithms. However, all the TEs were called either as present or absent while T-lex2 classifies the insertions as present, present/polymorphic, polymorphic, absent/polymorphic and absent.

### Statistical analysis

All the statistical analyses were performed using R (http://cran.r-project.org/). Sensitivity and specificity were estimated using individual PCR frequencies as we have the T-lex2 frequencies estimates using the same exact strains. Sensitivity and specificity were computed such as: (i) Sensitivity = number of correct presence calls / total number of presence calls in the validation dataset and (ii) Specificity = number of correct absence calls / total number of absence calls in the validation dataset. For the human dataset, the number of correct absence calls is not known because both polymorphic and absence calls are detected as absent in the validation dataset. Thus we used the positive predictive value (*i.e.,* number of correct presence calls based on T-lex2 / total number of presence calls) as a proxy for specificity and the false discovery rate (i.e., number of false presence calls based on T-lex2 / total number of presence calls) as proxy for the T-lex2 call precision.

## RESULTS

### T-lex2 is an improved and expanded version of T-lex

T-lex was originally designed to genotype and estimate frequencies of the known TE insertions in individual strain NGS data (26). The new version of T-lex, called T-lex2, incorporates numerous new features in order to improve genotyping and frequency estimation of known TE insertions (Table 1). T-lex2 assesses the quality of the calls by analyzing the genomic context of each TE insertion and detecting their TSDs that allows identifying miss-annotated TE insertions. T-lex2 takes into account the sequencing technology advances and can now handle PE sequencing and longer reads. Unlike the previous version, T-lex2 can be run for individual strains and for pooled NGS for the TE frequency estimates (Table 1).

### Accurate TE frequency estimates using NGS data from *D. melanogaster* individual strains

To test the quality of the T-lex2 calls in *D. melanogaster*, we compared T-lex2 TE frequency estimates with (i) frequency estimates previously obtained using a pooled-PCR approach (34,35,38) and (ii) frequency estimates obtained in this work by individual strain PCRs.

We first launched the pipeline to estimate the frequency of 755 TE insertions for which we have an estimate of their population frequencies based on a pooled-PCR approach (34,35,38). We run T-lex2 on 86 single DGRP strains (Supplementary Table S1; see Material and Methods). The TE call distribution, *i.e.* proportion of TEs called as present, absent, polymorphic, present/polymorphic, absent/polymorphic and no calls, is consistent among the strains, except for 14 strains that differ drastically (Supplementary Figure S1). For 11 out of the 14 strains, we observed more than 40% of no calls, and for the other three strains (RAL-379, RAL-362 and RAL-313), more than 70% of TE insertions were detected as present. As we suspect these 14 strains to be problematic (most likely a combination of poor sequence quality and extensive residual heterozygosity), we decided to remove them from our analysis and analyze the calls in the remaining 72 strains (Supplementary Figure S1).

Out of the 755 TE insertions, TE-analysis module detected six that were flanked by other TE insertions, and three that were part of segmental duplications (Supplementary Table S2). We removed these nine insertions because they are likely to produce wrong calls (see Material and Methods). We then filtered out seven TEs for which T-lex2 failed to return any calls, and 30 TE insertions for which T-lex2 fails to return a call for more than 75% of the strains (Supplementary Table S2). All of these removed TE insertions, 46 in total, are likely to be miss-annotated as we failed to call them in all or most of the strains. We thus estimated the population frequencies of 709 TE insertions using 72 strains. Note also that 65 out of the 709 TEs were identified in a first T-lex2 run as miss-annotated based on results of the TE-TSD detection module (see below). We manually re-annotated these 65 TE insertions, and we have re-run T-lex2 to obtain accurate frequency estimates of these TEs.

T-lex2 frequency estimates of the 709 TEs in 72 strains are significantly and positively correlated with previous estimates based on a pooled-PCR approach (Figure 5A: Spearman's *ρ* = 0.87, *P-val* ≪ 0.001). The vast majority of the very rare and fixed TE insertions are correctly detected by T-lex2 with 92.5% (357/386) of the very rare TE insertions absent and 85% (91/107) of the fixed TE insertions fixed in the population (Figure 5A). As the pooled-PCR and T-lex2 frequencies categories were not build the same way, we expected to observe intermediate profiles for the rare and common TE insertions. As expected, most of the rare TE insertions (114/146) are detected with a zero or low frequency, while most of the common TE insertions (67/70) appear frequent or fixed (Figure 5A).

**Figure 5.**
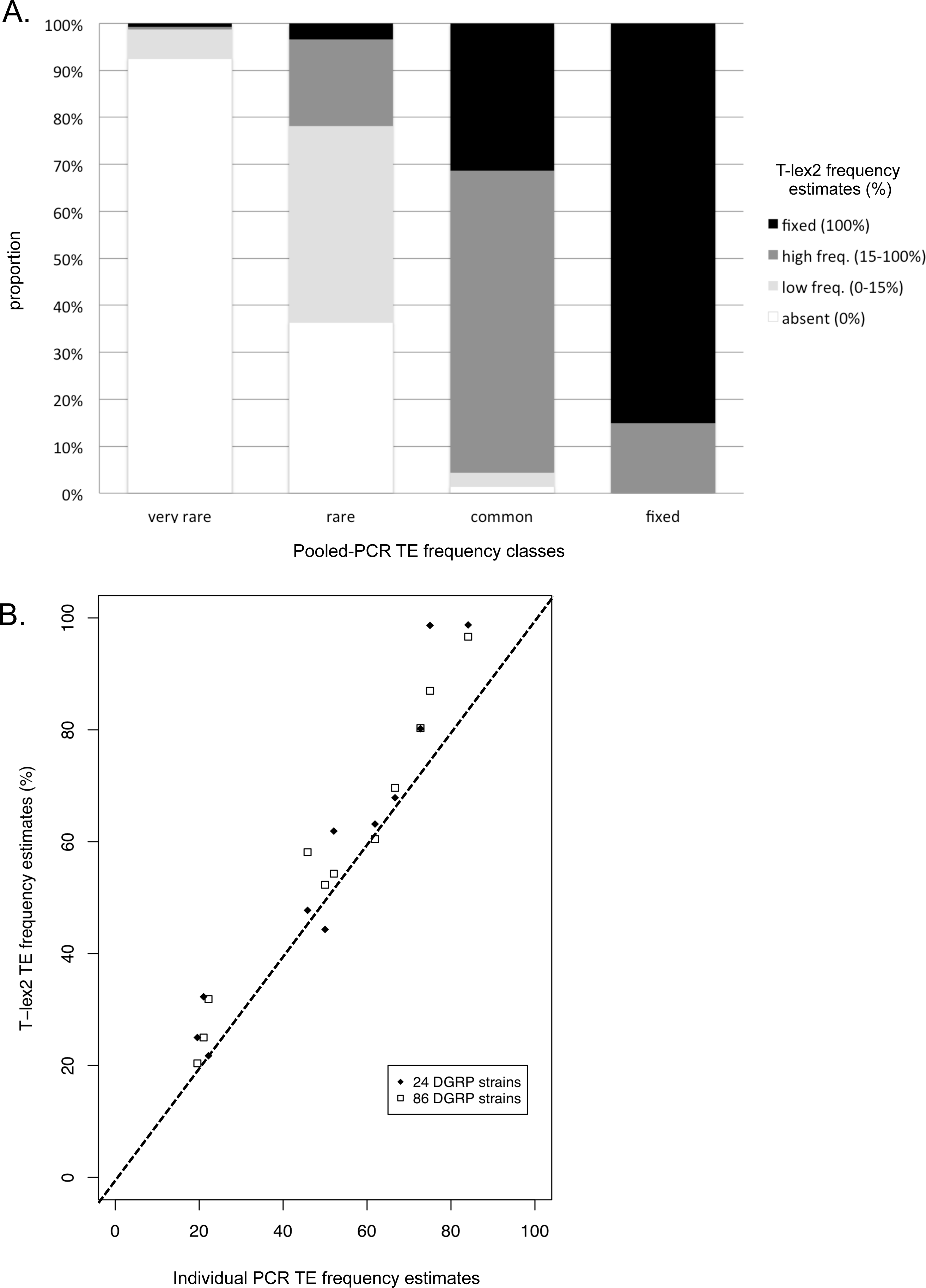
TE frequency estimates using *D. melanogaster* single strains. (A) Comparison of the frequency estimates of 709 TE insertions using T-lex2 and pooled PCR approach. (B) Frequency estimates of 11 TE insertions using T-lex2 and single PCR approach.

Overall, we identified only 1.5% (11/709) of clear discrepant estimates between pooled-PCR and T-lex2 frequency estimates. However, after manual curation, we suspected all these TE insertions to be miss-classified by the experimental approach or miss-annotated. For example, insertion *FBti0020042* is classified as present at very low frequency by T-lex2 but is detected at high frequency using the pooled-PCR approach. However, a single strain PCR approach confirms that this TE is present at a low frequency as estimated using T-lex2 (35). The reclassification of this TE insertion, allows to conclude that T-lex2 returns accurate TE frequencies with only ∼1% of wrong estimates.

To further test the accuracy of T-lex2 calls, we experimentally determined the frequency of 11 well-studied TE insertions known to be polymorphic and thus providing all types of calls (35). We then randomly selected 24 strains part of the bigger set of 86 DGRP strains (Supplementary Table S1) and performed single strain PCR for each of the 11 analyzed TEs. We experimentally called the 11 TE insertions in the same 24 DGRP strains (Supplementary Table S3). We first compared the frequency estimates for the 11 TE insertions obtained by T-lex2 using all the DGRP strains (86) and the subset of 24 strains. This analysis confirms the robustness of the frequency estimates using the subset of 24 strains (Figure 5B: Pearson's *ρ* = 0.96, *P-val* ≪ 0.001). Out of the 264 calls (11 detections in 24 strains), we removed 53 for which we failed to get a call experimentally or using T-lex2 and ended up analyzing 211 TE calls. We observed only 11 calls that were different between the experimental data and the T-lex2 calls (Supplementary Table S3). This represents 5.2% of putatively wrong calls that is close to the error rate previously estimated for the TE detection using PCR or the previous version of T-lex, around five percent in both cases (26,34). We finally conservatively estimated a high TE call accuracy with 99.14% of sensitivity and 89.58% of specificity (see Material and Methods). Thus, we overall observed accurate TE frequency estimates comparing the T-lex2 estimates with the pooled-PCR and the individual PCR experimental estimates.

### Accurate TE population frequency estimates using *D. melanogaster* pooled sequencing data

We called the presence/absence of the 709 selected TE insertions using data from a pool of 92 DGRP strains, containing the 86 single DGRP individual strains previously used (36) (Supplementary Table S1). For 29 out of the 709 selected TE insertions, T-lex2 failed to return a call using the pooled sample, most probably due to low read coverage in these particular regions of the genome. We thus ended up estimating the population frequency of 680 TE insertions (Supplementary Table S2). As expected, we found that the number of reads providing evidence for the absence is negatively correlated with the experimental TE frequency estimates (Figure 6A: Spearman's *ρ* = −0.60, *P-val* ≪ 0.001), while the number of reads providing evidence for the presence is positively correlated (Figure 6B: Spearman's *ρ* = 0.87, *P-val* ≪ 0.001). Interestingly, the number of reads supporting the presence does not seem to be strongly biased by the number of copies of TEs from the same family indicating the accuracy of our approach (see Material and Methods).

**Figure 6.**
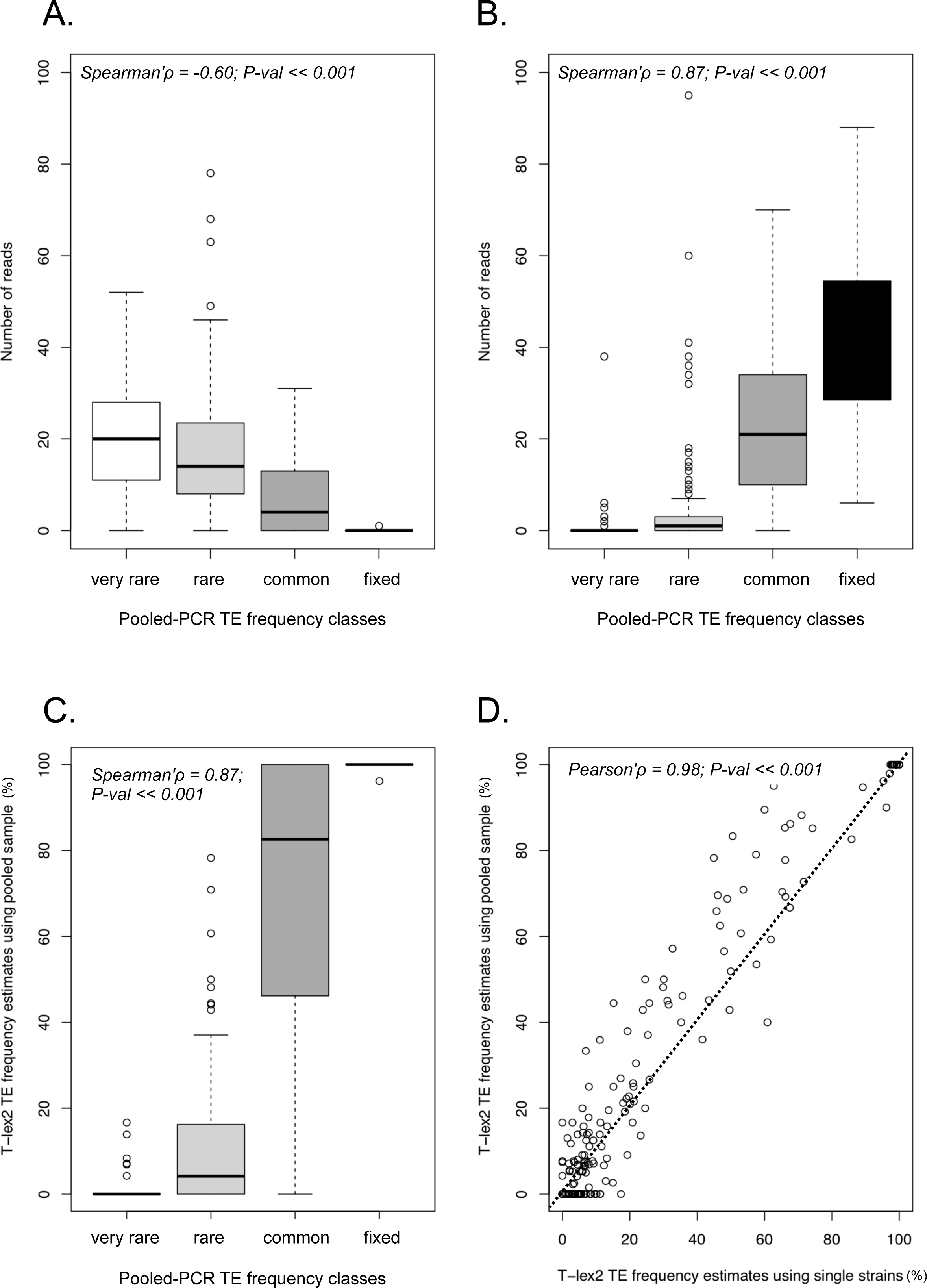
TE frequency estimates using *D. melanogaster* pooled samples. (A) Number of reads supporting the absence and (B) number of reads supporting the presence for each of the TE frequency classes. (C) T-lex2 frequency estimates using pooled sample *versus* TE frequency classes experimentally determined using a pooled-PCR approach. (D) Comparison of T-lex2 frequency estimates using single and pooled NGS data.

We observed significant and strong positive correlations between T-lex2 TE frequency estimates using the pooled sequencing data and the pooled-PCR determined frequencies (Figure 6C: Spearman's *ρ* = 0.87, *P-val* ≪ 0.001) and between the T-lex2 TE frequency estimates using the pooled sequencing data and the T-lex2 estimates using single strains (Figure 6D: Pearson's *ρ* = 0.98, *P-val* ≪ 0.001). This finding supports that the TE population frequency can be estimated with accuracy using pooled NGS sample only based on the read depth coverage.

For 25% of the TEs, all of them polymorphic, we noted an overestimation of the TE frequency using the pooled sample compared to the estimates using single strains (Figure 6D). However, for the majority of TEs the difference between the two estimates is <10%, which can be explained by the sampling effect when the pool and the individual strain libraries are constructed. To get accurate frequency estimates, each selected individual should be equally represented in the pooled library. Unfortunately, those conditions are hard to set up or control during library construction. For seven percent of the TEs, the discrepancy between the two estimates is >10% and the pool estimate is consistently higher than the individual strain estimate. However, we run T-lex2 in a different set of individual and pooled strains and although five percent of the TEs showed discrepant frequencies estimates between pooled and individual samples, these discrepancies were both over-and under-estimations (Supplementary Figure S2). These results suggest that T-lex2 frequency estimation is not biased and that the observed overestimation for the DGRP strains is likely due to some particular feature of this dataset such as sequencing coverage or number of strains used in pooled *vs* individual strains.

To assess the effect of the sampling and coverage on the TE frequency estimates using pooled data, we compared the TE frequency estimates using five different pooled datasets (Table 2). The read depth coverage varies from 20X to ∼60X (36), the number of strains from 42 to 92, and the number of flies sampled varies from 42 to 134. With only 42 flies pooled and a minimum coverage of 20X, the correlation is strongly positive (Table 2). The increase in the number of flies only does not seem to significantly improve the TE frequency estimates (Table 2). However, the increase of the coverage but not the number of strains improves significantly the TE frequency estimates (Table 2). Thus, the coverage matters more than the number of flies or strains pooled in the TE population frequency estimation using pooled sequencing samples. Read-depth coverage of 20X appears sufficient to obtain good estimates although the increase of the coverage allows reducing the number of errors in the calling.

**Table 2.**
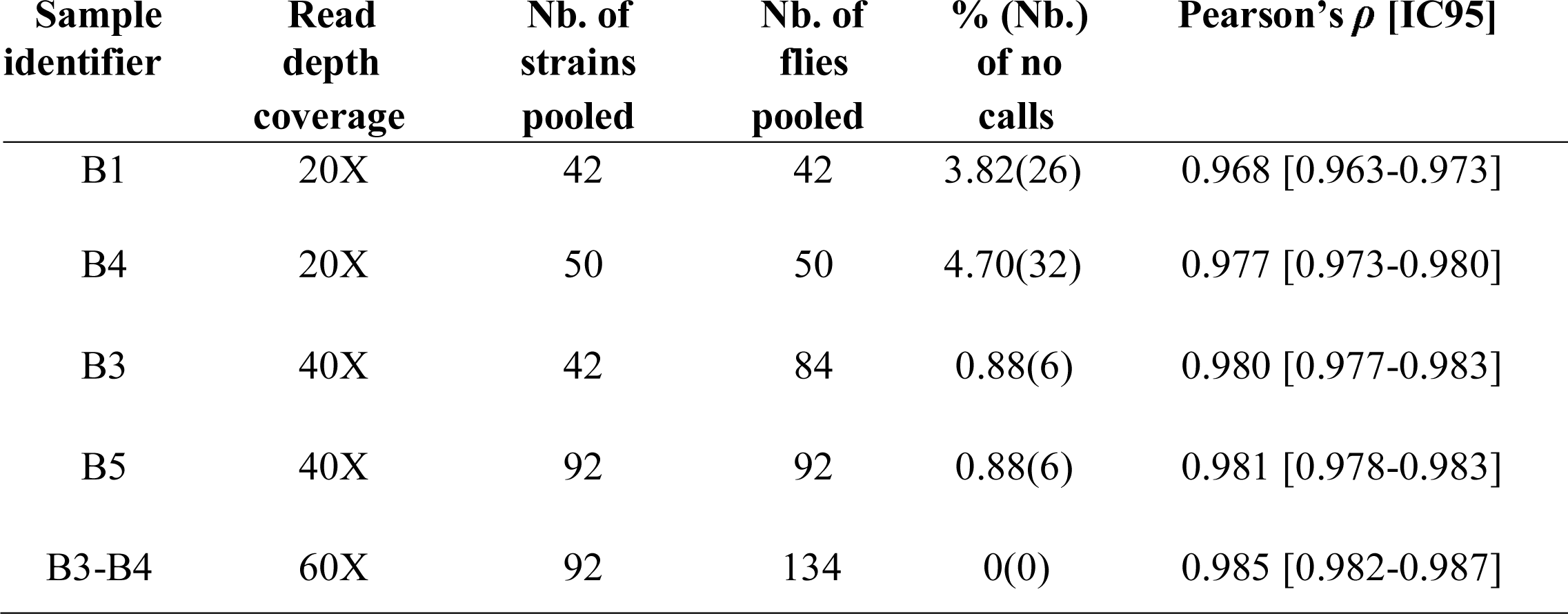
Sampling effect on the population frequency estimates of 680 TE insertions in *D. melanogaster*.

| **Sample identifier** | **Read depth coverage** | **Nb. of strains pooled** | **Nb. of flies pooled** | **% (Nb.) of no calls** | **Pearson's $\rho$ [IC95]** |
| --- | --- | --- | --- | --- | --- |
| B1 | 20X | 42 | 42 | 3.82(26) | 0.968 [0.963-0.973] |
| B4 | 20X | 50 | 50 | 4.70(32) | 0.977 [0.973-0.980] |
| B3 | 40X | 42 | 84 | 0.88(6) | 0.980 [0.977-0.983] |
| B5 | 40X | 92 | 92 | 0.88(6) | 0.981 [0.978-0.983] |
| B3-B4 | 60X | 92 | 134 | 0(0) | 0.985 [0.982-0.987] |

### T-lex2 detection of TSDs is unbiased and accurate

The new TE-TSD detection module of T-lex2 allows the automatic annotation of TSDs without *a priori* knowledge of the biology of the TEs. While only one copy of the TSD is present in the genome without the TE insertion (*i.e.*, the real TE absence), two TSD copies should be observed in the new constructed reference sequence used by T-lex2 for the absence detection (*i.e.*, the TE absence reference sequence constructed by computational removal of the TE sequence from the genome). Based on this expectation, the new T-lex2 module post-processes the results from the absence module to analyze the alignments of the reads spanning the TE insertion site to identify the TSDs. This new module looks for the expected missing sequence in the alignment flanking the TE insertion site (Figure 3). If the missing sequence corresponds strictly to the TSD, another copy of this TSD should be detected in the vicinity of the missing sequence. Contrary to most of the TSD detection approaches (39–41), no expected TSD length is required for T-lex2 TSD detection.

Out of the 587 TEs with at least one read providing evidence for the absence, T-lex2 detects TSDs for 390 of them (Table 3). TSD length ranges from two base pairs to 19 bp with an average and a median of five base pairs (Supplementary Figure S3). TSDs of the LTR retroelements are consistently short: four base pairs long on average. TSD lengths of DNA elements are also consistent but longer: seven base pairs on average. On the contrary, TSDs of the non-LTR retroelements (LINE) are not conserved within families and vary in length from four base pairs to 19 bp (Table 3). Overall, we identified for the first time TSDs for 10 TE families: six DNA transposon families and four non-LTR families (Table 3).

**Table 3.**
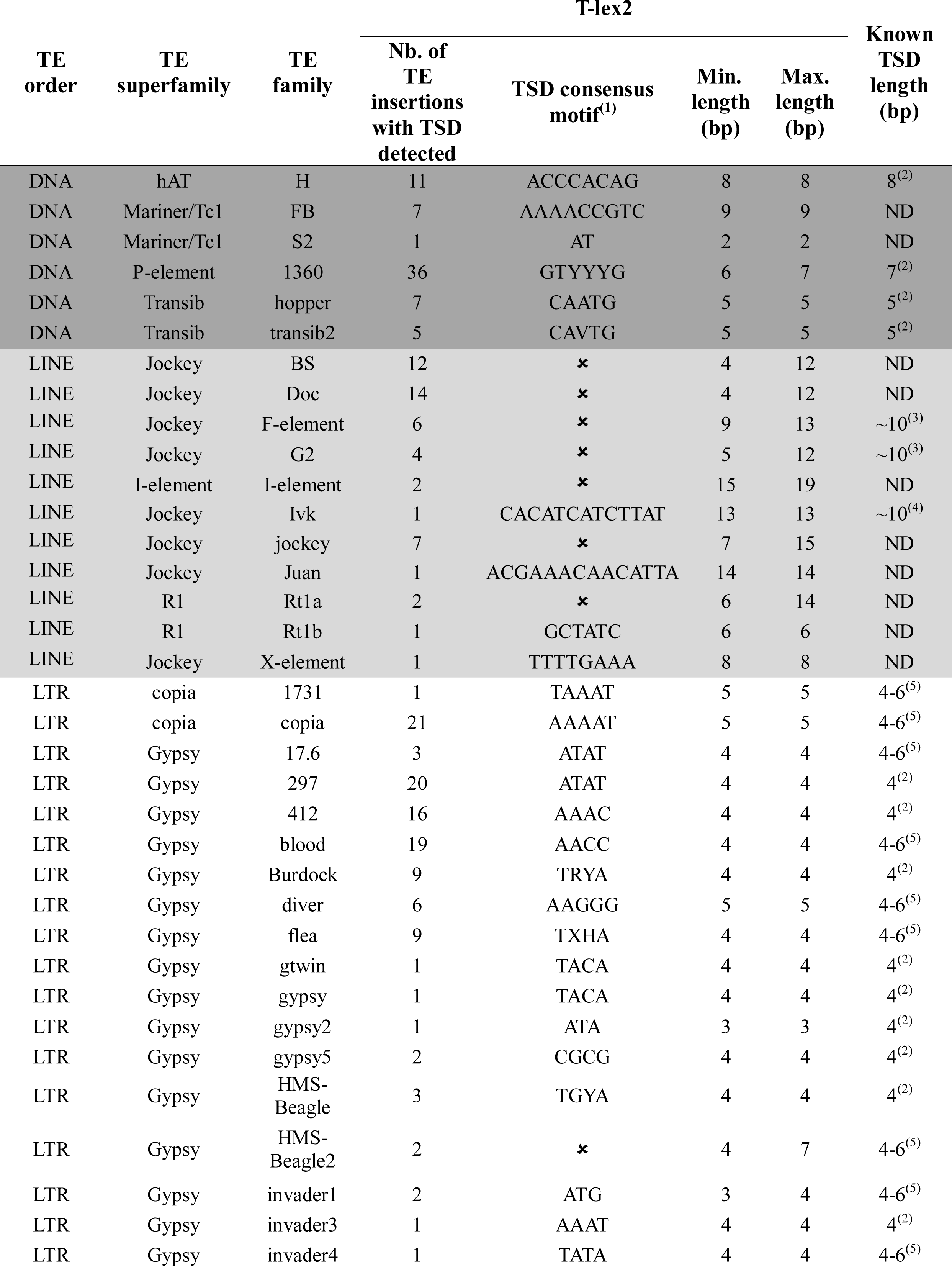
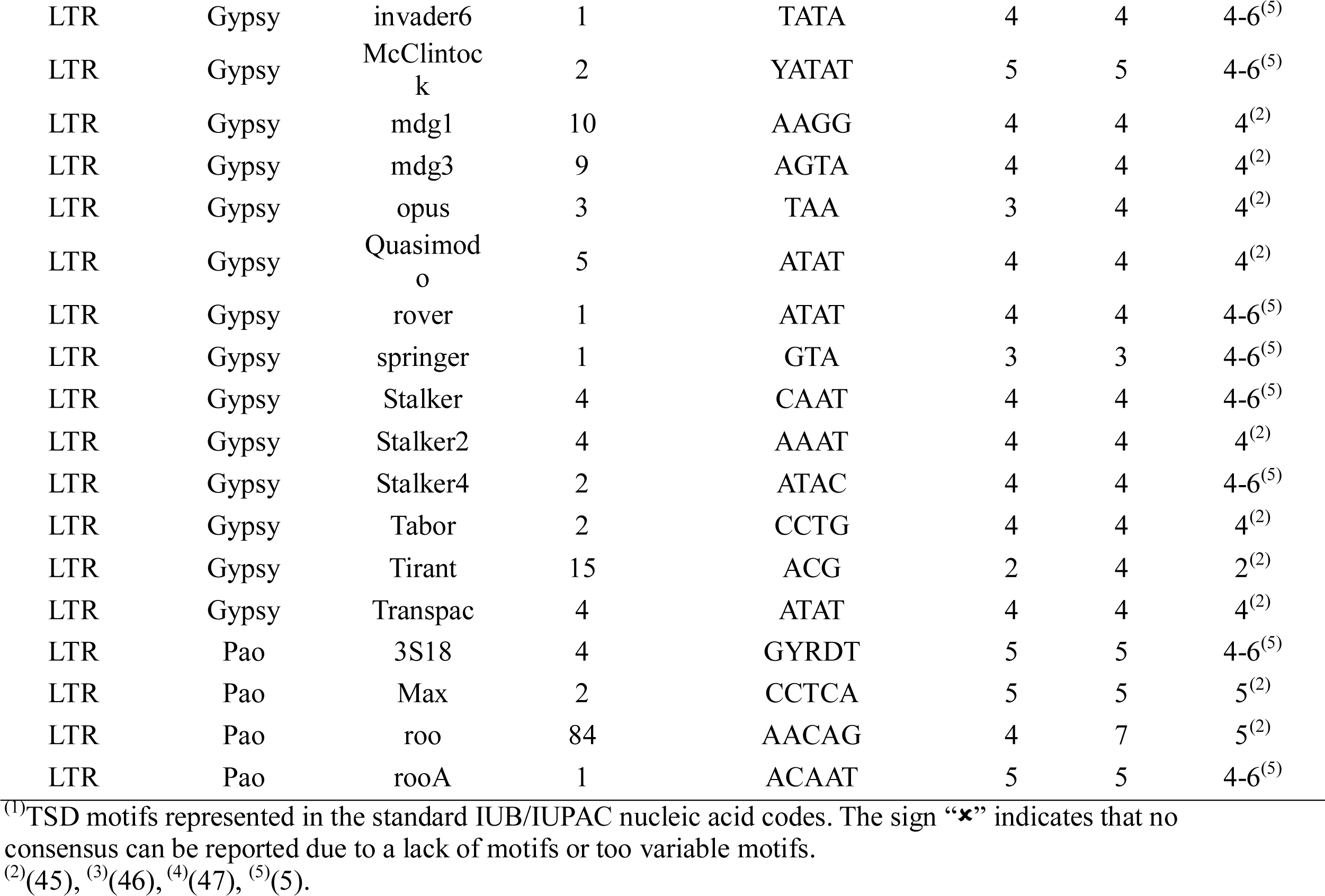
TSD detection for the 53 TE families analyzed in *D. melanogaster*. ND: Not determined.

To assess the quality of the TSD annotation using T-lex2, we compared 148 TSD results obtained by T-lex2 with the annotations obtained with *LTRharvest* software (39) (Supplementary Table S4). For 92 out of the 148 TE insertions, the TSD annotations are identical in length and motif. For seven TEs, TSDs have the same length but differ in motif (Supplementary Table S4) and for the other 49 TEs TSD motifs are identical but differ in length: seven TEs have shorter motifs and 42 TEs have longer motifs. All the longer TSDs correspond to shifts of only one base pair except for one TSD that is three base pairs longer than previously annotated. For 13 of these TEs, corresponding to *Copia* elements, T-lex2 results are supported by experimental studies indicating that *Copia* TSD are five base pairs long (42,43) and not four base pairs as detected with *LTRHarvest* (Supplementary Table S4) (39).

TE-TSD detection module did not identify TSDs for 197 TEs out of the 587 with at least one read providing evidence for the absence. TSD detection may fail for old TE insertions for which the TSD is too divergent, for truncated TEs, and when the boundaries of the TE insertions are not well annotated.

### T-lex2 allows re-annotation of TEs in *D. melanogaster*

Combining outputs from the TE-analysis module and the TE-TSD annotation, T-lex2 allows manual curation and re-annotation of TE insertions. To facilitate the identification of the putatively miss-annotated TEs, information on TEs for which the TSD detection failed and TEs with poly(A) tail longer than annotated, are also given in “Tfreq_combine” file. TSD detection fails for TEs that are longer (Figure 4A.II) and shorter (Figure 4A.III) than annotated. We thus manually curated the 197 TEs for which TSD could not be detected. For 65 out of the 197 TE insertions, manual curation confirms the miss-annotation. All these 65 TEs are detected experimentally and by T-lex2 at very low frequency (Supplementary Table S5). 64 of the 65 miss-annotated TE insertions are longer than annotated and they all correspond to non-LTR TE insertions: 33 have a longer poly(A) tail than annotated, and the other 32 TEs have one or the two junctions miss-annotated. The increase in length of these TEs ranges from five base pairs to 40 bp (12 bp on average; Supplementary Table S5). The only TE insertion detected as shorter than annotated is an LTR (*FBti0020118*; Supplementary Table S5). Thus, T-lex2 allows identifying miss-annotated TEs and manual curation allows re-annotating ∼10% (65/709) of the analyzed TEs.

### T-lex2 provides accurate TE presence/absence detection in human data

We also tested the accuracy of T-lex2 in the human genome, a genome with a very different TE composition and TE dynamics compared to that of the Drosophila genome. We used a curated subset of the Mobile Element Insertion (MEI) catalog (see Material and Methods) (37). Out of the 1,549 insertions analyzed, 755 TEs were excluded based on the results of the TE-analysis module because the repeat density in their flanking regions was higher than 50%. 12 additional TEs were removed from the final dataset because T-lex2 returned “no data” as a call. Thus, T-lex2 yielded highly accurate TE calls for a total of 782 TEs (Supplementary Table S6). We compared T-lex2 results with available validation data for the same subset of TEs obtained by PCR and a combination of different software (Table 4). T-lex2 results show an overall of 96,9% matches, which is higher than that obtained by the read pair (RP) and split read (SR) methodology used by Steward et al 2011(37) (93%). T-lex2 has a high TE call accuracy in human data with 97.65% sensitivity and 93.26% positive predictive value, a proxy for specificity (see Material and Methods). Finally, false discovery rate was low: 6.74% relative to the PCR and/or assembly validated calls from Steward et al. (2011) (37).

**Table 4.** Comparison between T-lex2 results and MEI catalog in humans. The number of matches between T-lex2 and MEI calls are given in parenthesis.

| T-lex2 | present | present/polymorphic | absent | polymorphic | absent/polymorphic |
| --- | --- | --- | --- | --- | --- |
|  | 267<br>(249) | 0<br>(0) | 279<br>(278) | 234<br>(229) | 2<br>(2) |
| MEI<br>catalog | present |  | absent |  |  |
|  | 255 |  | 527 |  |  |

## DISCUSSION

We present here the new version of the T-lex pipeline: T-lex2 (Figure 1). This integrated Perl computational pipeline designed to analyze TE insertions in NGS data is freely available and user-friendly since only one command line is necessary to run it (see http://petrov.stanford.edu/cgi-bin/Tlex.html or http://sourceforge.net/projects/tlex/). T-lex2 is also flexible. The pipeline is composed of five distinct modules that can be launched independently and each one of them includes a large number of options that allows performing flexible and customizable analyses (see T-lex2 manual at http://petrov.stanford.edu/cgi-bin/Tlex2_manual.html). T-lex2 runs with single-end and/or PE reads and can be customized for NGS datasets with different read lengths, read quality and read depth. Thus, T-lex is not only able to work with the different NGS datasets already available, but also with upcoming datasets resulting from improvements in the NGS technology.

Briefly, T-lex2 has improved and/or expanded the four modules present in T-lex and has incorporated a new module that allows detecting TSDs in an unbiased and accurate way (Table 1). Besides individual strain NGS data, T-lex2 can now estimate TE frequencies from pooled NGS data. This is an important new feature of T-lex2 since sequencing pooled samples is an inexpensive and efficient means of obtaining population genome data (25,36). Another important new feature of T-lex2 is the ability to identify TEs likely to return wrong calls *i.e.* miss-annotated TEs, TEs part of segmental duplications and TEs flanked by other TEs. This has been accomplished by the expansion of the TE-filter module of T-lex and by the incorporation of the TSD detection module (Figure 1). Being able to identify TEs likely to give wrong calls is especially important for the analyses of TE population frequencies in genomes that do not have a high-quality TE annotation. Note that even in an extremely high quality genome as the *D. melanogaster* one, we re-annotated ∼10% of the analyzed TEs. This result also supports the opportunities offered by the NGS data to improve TE annotations (44).

We tested the reliability of T-lex2 calls and frequency estimates using the fly and the human genome. Indeed, the pipeline can be used for any species for which NGS data and TE annotation is available. T-lex2 can be run for all type of TEs in any species because (i) it is a flexible pipeline that allows to customize runs for NGS datasets of different qualities, (ii) it is able to identify putatively wrong calls and re-annotate miss-annotate TEs, and (iii) no information about the biology of the TE is needed to get accurate frequency estimates.

There are other available tools that can be used to estimate TE frequencies (10,21–25). However T-lex2 is to date the only tool that (i) combines presence and absence detection analysis that allows the identification of heterozygotes and/or polymorphic TE insertions; (ii) can be used for individual and pooled samples; (iii) provides the user with an output file with the frequency of the analyzed TEs; (iv) can be easily customize to be run with different NGS datasets in any organism; and (v) assess the quality of the calls allowing to re-annotate miss-annotated TEs. Thus overall, T-lex2 is the most broadly applicable and flexible tool available to date.

The flexibility of T-lex2, exemplified by the incorporation of the TE-TSD detection module in this new version, will allow us to easily add new modules in a near future. We are currently working on a new module designed to identify TEs not annotated in reference genomes. However, T-lex2 can be used to genotype and estimate frequencies of *de novo* TEs identified with already available software such as RetroSeq (22). The ability of T-lex2 to re-annotate TEs would be specially useful to study this *de novo* TEs as their junctions might not be as well annotated as the ones in the reference genomes.

Overall, we are providing a versatile tool that allows exploring the impact of TEs on genome evolution as well as learning about TE biology. By analyzing the frequency of TEs in different populations, we will be able to determine which proportion of the TE-induced mutations are subject to strong purifying selection or are likely to be evolving under positive selection. Additionally, accurate annotation of TE insertions will allow us to study for example the target site preferences of different TE families. The availability of a tool such as T-lex2 improves our ability to explore TE dynamics and TE biology in an increasing number of species. This is important, as we cannot hope to understand genome structure and function without a thorough understanding of its most active, diverse, and ancient component.

## ACKNOWLEDGMENTS

We would like to thanks the ScaleGenomics (http://scalegen.com/) and Proclus (Stanford University; http://www.stanford.edu/) computational platforms that we used to investigate this study.

